# Statistical Analysis of Three-Dimensional Chromatin Packing Domains Determined by Chromatin Scanning Transmission Electron Microscopy (ChromSTEM)

**DOI:** 10.1101/2021.05.24.445538

**Authors:** Yue Li, Vasundhara Agrawal, Ranya K. A. Virk, Eric Roth, Wing Shun Li, Adam Eshein, Jane Frederick, Kai Huang, Luay Almassalha, Reiner Bleher, Marcelo A. Carignano, Igal Szleifer, Vinayak P. Dravid, Vadim Backman

## Abstract

Chromatin organization over a wide range of length scales plays a critical role in the regulation of transcription and deciphering the interplay of these processes requires high-resolution, three-dimensional, quantitative imaging of chromatin structure *in vitro*. Herein, we introduce ChromSTEM, a method that utilizes high angle annular dark-field imaging and tomography in scanning transmission electron microscopy combined with DNA-specific staining for electron microscopy. We utilized ChromSTEM to quantify chromatin structure in cultured cells and the statistical packing behavior of the chromatin polymer. Using chromatin mass and density analysis, we observed that chromatin forms spatially well-defined higher-order domains which are around 100 nm in radius, with a radially decreasing mass-density from the center to the periphery. Although the morphological properties of the domains vary within the same cell line, they seem to exhibit greater heterogeneity across cell lines, underlying a potential role of statistical chromatin packing in regulating cell-type-specific gene expression.

## Introduction

Three-dimensional chromatin packing in the cell nucleus plays an important role in regulating numerous cellular processes, and large-scale alterations in chromatin structure are associated with cancer, neurological and autoimmune disorders, and other complex diseases (*1–3*). The fundamental repeating unit of chromatin is the nucleosome, in which 147 bp of the DNA is wrapped around a core histone octamer (*4*). The core particle adopts a squat cylindrical shape, with a diameter and height of approximately 11 nm and 5.5 nm, respectively (*5*). The nucleosome is the first level of higher-order packing of the chromosomal DNA. Nucleosomes are connected by linker DNA, which altogether form what is referred to as the 10-nm chromatin fiber (*6*). Central to the textbook view of chromatin packing is that 10-nm chromatin fibers assemble into 30-nm fibers, that further fold into 120-nm chromonema, to 300- to 700-nm chromatids, and ultimately, mitotic chromosomes (*7–10*).

However, the key tenant of this view, the 30-nm fiber, has been challenged by an abundance of recent evidence. Various studies using cryo-electron microscopy, small-angle X-ray scattering, electron spectroscopy imaging, and super-resolution microscopy failed to observe 30-nm fibers in interphase chromatin or mitotic chromosomes in numerous cell lines (*11–14*). For example, Ricci et al. observed the existence of heterogeneous nucleosome ‘clutches’ at the level of the primary fiber, the size of which depend on epigenetic state and cell type (*14*). Recently, a combination of DNA-specific staining (ChromEM) and multi-tilt electron tomography (ChromEMT) observed *in situ* that the chromatin fiber consists of disordered chains that have diameters between 5 to 24 nm during both interphase and mitosis, with a higher packing concentration in mitotic chromosomes (*15*). Altogether, these studies suggest that the chromosome organization is constructed by 10-nm fibers without folding into 30-nm fibers (*12, 16, 17*). In this new paradigm, the 10-nm fibers condense into highly disordered and interdigitated states, which may be constantly moving and rearranging at the local level (*18–20*).

Despite their dynamic and fluid-like nature, several complementary imaging studies have revealed higher-order, domain-like structures above the level of the primary fiber. ‘Chromomeres’, punctate chromatin particles around 200-300 nm in diameter, have been observed in both interphase chromatin and mitotic chromosomes using stimulated emission depletion (STED) microscopy (*21*). A recent study employing photoactivated localization microscopy (PALM) live-cell imaging in mammalian cells determined that nucleosomes are arranged in physically compact chromatin domains with a diameter of around 160 nm (*22*). The dynamics of these chromatin domains were correlated with replication domains, which other super-resolution and electron microscopy studies have determined range in diameter between 110-150 nm (*22–25*). 3D-structured illumination microscopy (SIM) imaging demonstrated that DNA labeled with fluorescent *in situ* hybridization (FISH) forms chromatin domain clusters (CDCs) of around 120 to 150 nm in diameter with radially arranged layers of increasing chromatin compaction from the periphery towards the CDC core for mammalian cells (*26*).

Meanwhile, chromatin conformation capture (3C) and related methods (4C, 5C, Hi-C, Dip-C) have revealed that the eukaryotic genome is partitioned into topologically associating domains (TADs) at the scale of several hundreds of kilobases (kbs) and smaller loop domains (*27–30*). Recently, high-resolution imaging experiments have visualized TADs identified by Hi-C as domains in single cells, providing a link between the nanoscopic spatial structures and genomic domains (*31, 32*). Combining both *in vivo* and *in vitro* data, such domain structure may represent fundamental building blocks used to assemble higher-order compartments, and ultimately an interphase chromosome partitioned into euchromatin and heterochromatin. Additionally, these higher-order chromatin structures potentially play an important role in DNA-based processes, such as transcription, replication, and repair, and perhaps extends to complex processes, such as aging and diseases like cancer (*33–36*). However, a precise understanding of the internal structure of these chromatin domains is currently lacking, and new experiments with higher resolution, such as ChromEMT, are needed.

In parallel with experimental findings, many polymer models have been proposed to understand the statistical folding of these chromatin domains. A fractal globule model that describes chromatin as a collapsed polymer where topological constraints result in a hierarchy of non-entangled structures, explains earlier Hi-C results but was later challenged by data with higher resolution (*37, 38*). Additionally, chromatin domains observed by recent PALM imaging adopt a folding deviation from the fractal globule model at large length scales (*39*). More recently, a logarithmic fractal model was proposed to describe the large-scale organization of chromatin based on small-angle-neutron-scattering (SANS) experiments (*40*). Additional statistical models of chromatin have been proposed, including the novel self-returning random walk (SRRW) model, which depicts chromatin as non-globular, porous, and irregular “tree” domains and is able to reproduce key experimental observations including TAD-like features observed in Hi-C contact maps (*41*).

However, previous imaging and analysis methodologies are insufficient to determine the underlying principles of chromatin organization and function. As the physical entity of DNA, the chromatin higher-order structure can modulate the kinetics and efficiency of transcription reactions within domains with similar statistical packing behavior (e.g. chromatin packing scaling), influencing the regulation of global transcriptional patterns, and consequentially, the phenotypic plasticity of a cell (*42–45*). To properly characterize this statistical packing behavior of domains requires an imaging modality that provides structural data with high resolution at the level of the DNA base pair, the functional unit of transcription. Employing such structural data, we could further explore the assembly, packaging, and morphology of previously observed chromatin domains *in situ* to understand their functional significance by employing polymer physics-based analysis methods.

In this paper, we utilized scanning transmission electron microscopy tomography with ChromEM staining (ChromSTEM) to resolve the 3D chromatin organization for two mammalian cell lines *in vitro.* Importantly, ChromSTEM provides visualization of DNA structure down to sub-3nm-resolution with images intensity directly correlated with DNA density. We observed that chromatin fibers fold into distinct, anisotropic packing domains in which the mass scaling follows a near-power-law relationship. We further quantified the physical properties related to material transportation and gene accessibility of these domains, including chromatin volume concentration (CVC) and exposure ratio. Finally, we revealed such properties can be predicted by chromatin packing scaling and domain size, unveiling a potentially important link between the chromatin structure and functionality.

## Results

### ChromSTEM imaging of chromatin organization in mammalian cells

Following the ChromEM protocol reported previously, we labeled the DNA of human pulmonary adenocarcinoma epithelial (A549, Fig. 1, Mov. S1) and human fibroblast (BJ, Fig. S1, Mov S2) cells to characterize the chromatin packing behavior in two genetically distinct cell lines. After resin embedding, the labeled regions can be identified based on image contrast in bright field optical micrographs: the photo-oxidized cells appeared significantly darker than the non-photobleached cells (Fig. 1A-C). Dual-tilt STEM tomography in HAADF mode was performed for part of the nucleus where a hetero/euchromatin interface was observed on a 100 nm resin section (Fig. 1D). Unlike the near-binary image contrast from the conventional EM staining method (*15*), ChromSTEM provides continuous variations of the DNA contrast inside the nucleus. Each tilt series was aligned with fiducial markers in IMOD and reconstructed by a penalized maximum likelihood (PLM-hybrid) algorithm in Tomopy (*46, 47*). The two sets of tomograms were combined in IMOD to suppress missing cone (Fig. 1E) artifacts (*48*). The final tomography (Fig. 1F) has a nominal voxel size of 2.9 nm, with clearly resolved nucleosomes (Fig. 1G) and linker DNA (Fig. 1H). We also identified several distinct higher-order supranucleosomal structures, such as stacks and rings (Fig. 1I-1J). Examples of the full stack of tomography are shown in Mov. S1-S3. We also rendered the 3D volume of the chromatin in the volume viewer in FIJI (Fig. 1K-1L, Mov. S4-S9). The voxel intensity of the tomogram was used for color-coding (*49*).

**Fig.1.**
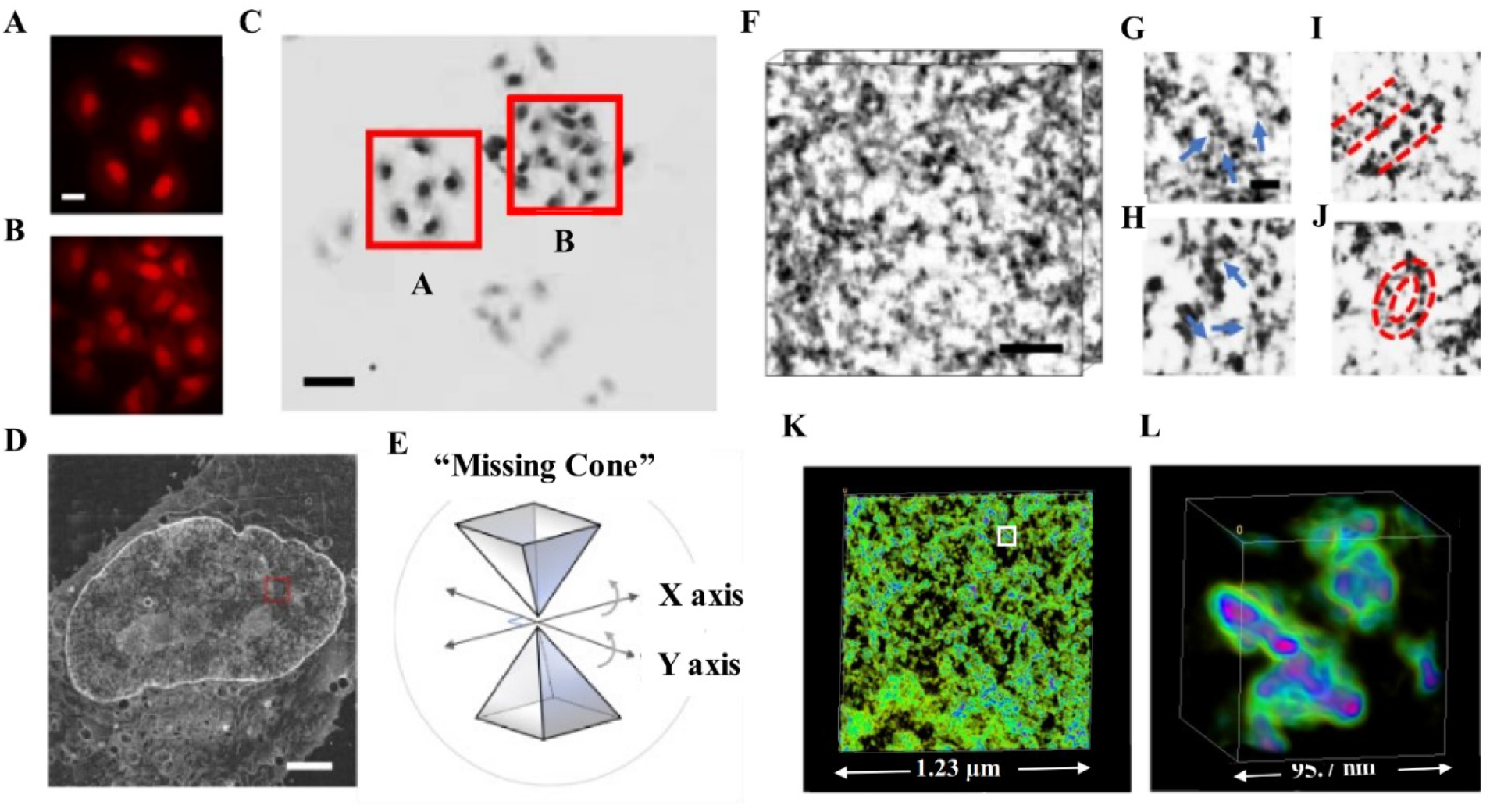
ChromSTEM tomography reconstruction of chromatin in an A549 cell. (**A-B**) The DRAQ5 photo-oxidation process takes 7 min for each region of interest. Scale bar: 10 μm. (**C**) The labeled regions were more intensely stained than the nearby regions (red squares; the letter corresponds to the regions in the left panels). Scale bar: 20 μm. (**D**) STEM image of a 100 nm thick section of an A549 cell in HAADF mode. Scale bar: 2 μm. (**E**) Schematics for dual-tilt tomography. The sample was tilted from −60° to 60° with 2° increments on two perpendicular axes. (**F**) 3D tomography of the A549 chromatin. Scale bar: 120 nm. (**G-H**) The fine structure of the chromatin chain: Nucleosomes (blue arrows in **G**), linker DNA (blue arrows in **H**) supranucleosomal stack (red dashed lines in **I**), and ring (red dashed circles in **J**). Scale bar: 30 nm. (**K-L**) 3D rendering of the chromatin organization, the pseudo-color was based on the intensity of the tomograms (Mov. **S4-S6**). (**L**) A magnified view of the region labeled by a white square in **K**. In **L**, pink and green regions represent high and low DNA density regions, respectively.

### ChromSTEM reveals chromatin packing domains with mass scaling behavior

Due to its semi-flexible nature, the chromatin polymer can, in principle, adopt an infinite number of potential 3D conformations which are not conserved temporally or across cell populations (*50*). However, the statistical properties of the chromatin polymer can be characterized, and are predicted to obey scaling laws, which describe how the number of monomers or, equivalently, the mass of the polymer, varies with the size of the physical space it occupies *(51)*. Depending on the balance of the free energy of polymer-polymer compared to polymer-solvent interactions, under dilute, equilibrium conditions a homopolymer chain, where all monomers interact in the same way, is expected to exhibit mass scaling characterized by a power-law relationship between the mass (*M*) and the size *r* at certain length scales: *M* ∝ *r^D^*, where *D* is the packing scaling of the polymer. When the interaction of monomers with the solvent is preferred, 5/3 < *D* < 2. When self-interaction is preferred, the polymer will collapse and adopt a scaling between 2 and 3. In a good solvent, *D* = 5/3 and the polymer adopts a self-avoiding random walk. When a polymer’s self-interaction and interaction with the encompassing solvent are equally preferred, as in the case of an ideal chain in a theta solvent, *D* = 2. A special case of *D* = 3 is the fractal globule structure. Importantly, in heteropolymer systems, when multiple monomer types alternate in sequential blocks along the linear polymer chain, for length scales above the size of the individual domains formed by blocks, *D* = 3 can also indicate a random distribution of spatially uncorrelated domains.

However, chromatin exists as a heteropolymer with the monomers, i.e., nucleosomes, possessing varying biochemical properties in the form of DNA modifications such as methylation, and those associated with post-translational histone modifications. Chromatin conformation can be further influenced by active molecular mechanisms that impose additional topological constraints, including CTCF-cohesin- or transcription-dependent looping, interactions with nuclear lamins, and phase separation driven by chromatin-associated proteins such as HP1 (*52, 53*). Therefore, at any given point in time, chromatin conformation is determined by such active constraints in addition to the balance between chromatin-chromatin and chromatin-nucleoplasm interactions, resulting in a non-equilibrium system. Additionally, chromatin occupies a significant volume fraction within the nucleus. As a result of such non-dilute conditions, the rules of polymer physics does not guarantee that the entire chromatin system can be described using the same power law-scaling relationship. In other words, there may be separate regimes with different mass scaling behavior. In general, different scaling behavior for the chromatin polymer system may exist at different length scales because (1) the primary chromatin chain may exhibit different intra-chain scaling, and (2) within certain scaling regimes that define domains with similar statistical packing properties, individual domains may be characterized by different values of *D* or the size of the individual domains may vary, which would alter the limits of this power-law scaling regime.

To elucidate the chromatin structure within the cell nucleus, we investigated the mass scaling of the binary masks of chromatin segmented from the tomograms (Fig. 2A-D). In the analysis, we treat the heterogeneous chromatin fibers, as reported by Ou et al. using ChromEMT, to be the fundamental element in building higher-order structures (*15*). The details of the segmentation procedure can be found in Fig. S2A. Practically, the 3D mass scaling relationship is defined as how the total amount of chromatin (*M*) enclosed within a volume 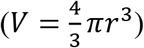 changes with its radius *r*. The 2D case can be described as a slice of the 3D system by a horizontal plane. In this case, *M* is the amount of chromatin enclosed within an area (*A* = *πr*^2^). The derivative of the area results in the perimeter, which represents the 1D case. Therefore, in the 1D scenario, *M* is the amount of chromatin positioned on the circumference of a circle (*P = 2πr*). We refer to the 1D case as “ring mass scaling”. We calculated the ring, 2D and 3D mass scaling by performing linear regression in the log-log scale on the mass scaling curves for the given dimensions. Since the packing scaling for different dimensions can be approximated from each other for the same fractals structure by the law of additivity of fractal codimensions (*54*), we confirmed from our calculations that the 3D mass scaling exponent can be estimated using the 2D and ring mass scaling (supplementary methods, Fig. S3A).

**Fig.2.**
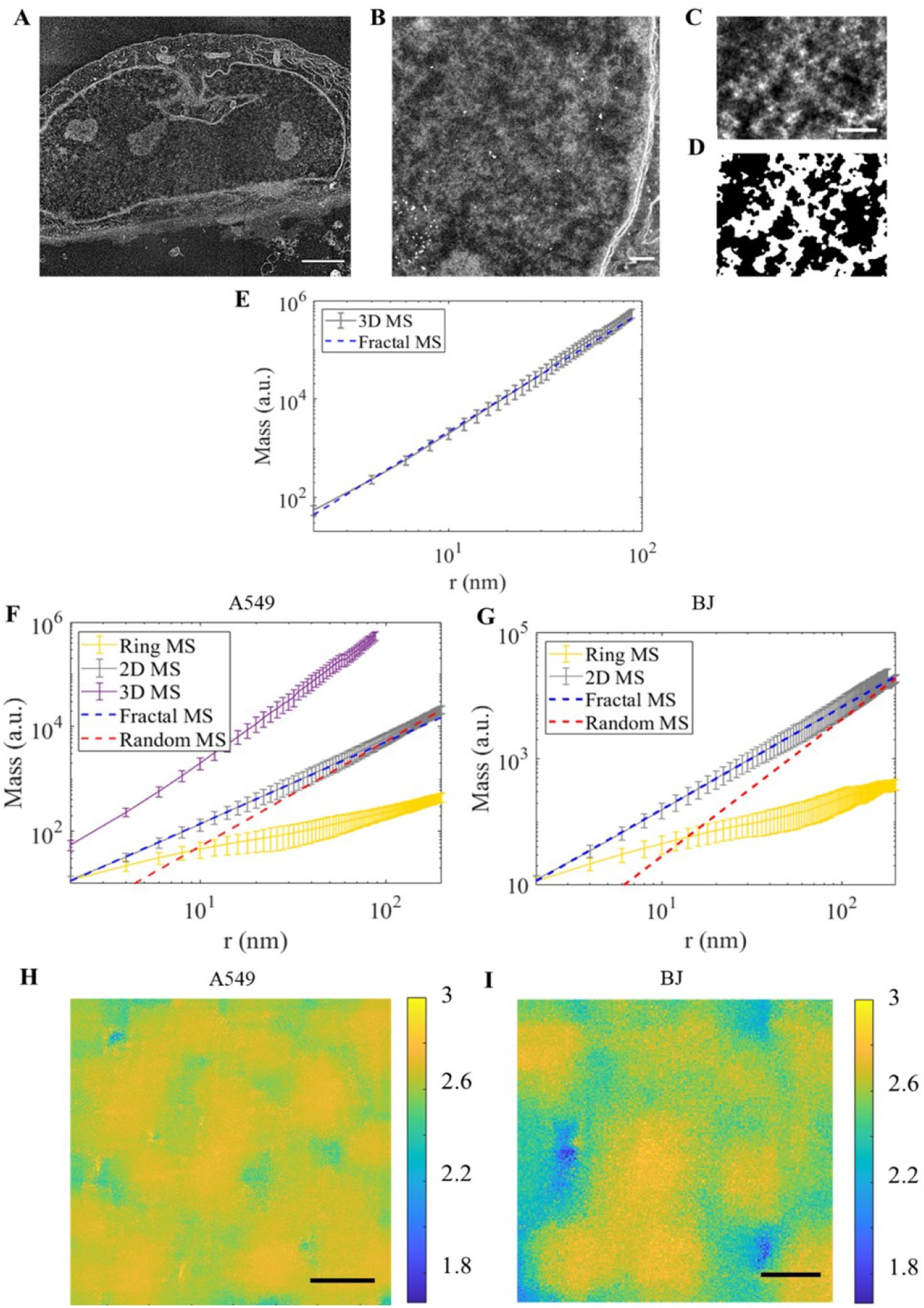
Chromatin folds into packing domains which have similar mass scaling behavior. (**A**) STEM HAADF image of a 150 nm section of a BJ cell nucleus for tomography reconstruction. Scale bar: 2 μm. (**B**) A magnified view of the chromatin in the nuclear periphery of the same cell in **A** with gold fiducial markers. The intensity variation of the image shows that the chromatin packs at different densities throughout. Scale bar: 200 nm. (**C**) A virtual 2D slice of the chromatin of a BJ cell after tomography reconstruction. Scale bar: 100 nm. (**D**) Binary mask of the chromatin location for the same area in **C**. The mass scaling analysis was performed on the binary masks. (**E-G**) The average mass scaling (MS) curve at different dimensions of the chromatin was imaged from four A549 (**E, F**) and four BJ (**G**) cells. The mass scaling analysis was conducted with randomly selected centers within the field of view, and a mean mass scaling curve is shown for each dimension, the error bar represents the standard deviation. **(E)** 3D mass scaling curve exhibits power-law behavior with a single scaling up to *r* = 90 nm. Slope, *D* = 2.59 ± 0.02 was obtained from linear regression from *r* = 2 nm to *r* = 90 nm. We found that the 3D mass scaling exponent can be approximated using the 2D case and the 1D case: *D*_3*D*_ = *D*_2*D*_ + 1, and *D*_3*D*_ = *D*_1*D*_ + 2. Two regimes of mass scaling with different packing scaling *D* can be identified. In the 2D cases for both A549 cells **(F)** and BJ cells **(G)**, the MS curve starts with a packing scaling with *D* < 3 (blue dashed line) and smoothly transitions to values close to *D* = 3 (red dashed line). (**H**) Spatial distribution of packing scaling *D* within the domain regime of an A549 cell. The color represents the value of *D*. Scale bar: 200 nm. (**I**) The spatial distribution of *D* within the domain regime of a BJ cell. Scale bar: 100 nm.

As ChromSTEM provides only a snapshot of the chromatin conformation at a single time point, we randomly sampled different regions within the field of view and calculated the mean mass scaling to capture the statistical behavior. For four A549 cells with a total volume of 1.16 μm^3^ resolved at a voxel resolution of 2.0 to 2.9 nm, we obtained the mass scaling curves at all three dimensions (Fig. 2E, F). A total volume of 0.09 μm^3^ was reconstructed from four BJ cells at a nominal voxel resolution of 1.8 to 2 nm and mass scaling analysis was performed (Fig. 2G). In order to obtain the packing scaling and identify length scales where a single scaling exponent cannot sufficiently describe the packing behavior, we evaluated the derivative of the log-log scale of the 3D and 2D mass scaling curves as a function of *r*. The slope, *D_log_* was defined as a linear regression fit to the log-log scale of the mass scaling curves that has an error of less than 5%. This linear regression fit, *D_log_* should be equivalent to the packing scaling, *D* within the power-law scaling regime, which we define when length scales associated with *D_log_* extend across at least one order of magnitude. From our 3D mass scaling analysis on A549 cells, we observed a power-law mass scaling regime extending from 2 nm to 90 nm with a fitting parameter of *D_log_* = 2.59 ± 0.02 (Fig. 2E, blue dashed line). However, because of the maximum section thickness of 180 nm for our A549 cells, our 3D analysis was unable to evaluate potential mass scaling behavior above 100 nm. Additionally, we did not perform the 3D mass scaling analysis for BJ cells, as the thickness of the reconstructed section of BJ cells is smaller than 70 nm, and the 3D mass scaling curve would only extend up to 35 nm.

Due to the intrinsic length-scale limitation of 3D mass scaling because of limited section thickness, we then performed the mass scaling analysis at different dimensions for both A549 and BJ cells. Employing the law of additivity of fractal codimensions, we calculated the 3D mass scaling exponent from 2D and 1D mass scaling curves as (Fig. S3B): *D*_3*D*_ = *D*_2*D*_ + 1, and *D*_3*D*_ = *D*_1*D*_ + 2 (*54*). For both A549 and BJ cells, we first evaluated the slope of the 2D mass scaling curve in the log-log scale along its entire length using a 12 nm sliding window. By estimating the local slope for small ranges of *r* along the entire length of the 2D mass scaling curves, two distinct regimes were identified. The first regime stretched up to *r* e~ 90 nm, followed by a gradual increase in the local log-log derivative towards a value of 3. Similar to the 3D mass scaling analysis, for A549 cells (Fig. 2F), we then obtained the slope of linear regression, *D_log_* = 2.60 ± 0.01 for 2 nm < *r* < 88 nm (blue dashed line). Above these length scales (*r* ~ 90 nm), the slope continuously increases until it approaches 3 for *r* > 252 (red dashed line) up to 300 nm. Similarly, for BJ cells (Fig. 2G), the fitting parameter for the linear regression was estimated to be *D_log_* = 2.62 ± 0.01 (blue dashed line) for 2 nm < *r* < 86 nm, and *D_log_* approaches 3 (red dashed line) for *r* > 182 nm. The shift from the domain regime to the supra-domain regime (*D_log_* ~ 3) is continuous, as opposed to a sharp, biphasic transition. The implications of this result on the conformation of chromatin within packing domains and a detailed investigation of the boundary of the domain regime will be discussed later.

In addition, the ring mass scaling curve exhibits a third regime from 2 nm < r < 10 nm for both cell lines (Fig. S3C-D), which can be interpreted as the chromatin chain regime. The upper length scale (10 nm in radius) agrees with the upper limit of the primary chromatin chain size (24 nm maximum diameter) (*15*). However, this regime is elusive on the mass scaling curves of higher dimensions, possibly caused by limited tomography resolution.

Therefore, both the 3D and 2D mass scaling analyses suggest that for length scales up to 90 nm, chromatin packing domains with similar mass scaling properties internally. From 2D mass scaling analyses, at larger supra-domain length scales, a gradual increase in *D* towards a value of 3 can potentially be explained by either a variability of packing domain sizes or by an overlap between domains that are being averaged out in the mass scaling analysis. To test these hypotheses, we mapped the spatial distribution of packing scaling *D* for the domain regime from 2D mass scaling curves calculated within a moving window (300 pixels x 300 pixels) on each virtual 2D tomogram for each cell. We then evaluated the radial density and mass scaling profile of the identified domains as a function of distance from the center of the domains. Overlap between neighboring individual domains can be identified by a gradual increase in *D_log_* from the *D* within domains to *D_log_~3* within the overlapping region. For both A549 (Fig. 2H) and BJ (Fig. 2I) cells, we observed domain-like structures with the mass scaling within the domains following a near power-law relationship with *D_log_* < 3 leading to a sharp transition to the supra-domain regime(with *D_log_* ~ 3. Altogether, this suggests that chromatin folds into spatially separable packing domains all of which have similar mass scaling behavior within domains but differ in their genomic and physical sizes as well as the value of the mass density scaling.

### Quantifying domain size and chromatin packing behavior at the domain boundary

Our previous analysis averaged mass scaling behavior from all domains analyzed within a given field of view. Next, we wanted to better characterize the mass-scaling behavior of individual domains. Here, we interpret length scales in our mass scaling analysis as the physical distance from a “domain center region”, areas of higher *D* at the center of domains. At small length scales within this “center region”, chromatin has similar mass scaling behavior. The boundary of this scaling behavior, equivalent to the size of the domain, can be defined as the length scale where the statistical behavior, such as the mass scaling, of the chromatin deviates significantly from the statistical behavior of the chromatin within the “center region”. At the same time, for an isolated domain with *D* < 3, the chromatin density decreases from the “center region” to the periphery. For spatially separable domains which exhibit distinct mass scaling behavior, the radial chromatin density per domain is expected to initially decrease, followed by a recovery due to the intersection with other domains. Thus, the boundary of a single domain can also be dictated by the radial chromatin density profile distribution.

To begin this more detailed analysis, we first identified the “domain center region” of each packing domain. From the spatial distribution of the packing scaling (Fig. 3A), we applied Gaussian filtering and local contrast enhancement before segmentation (Fig. 3B). Regions with the top 10% of the elevated *D* values were included as the “domain center region” (green areas in Fig. 3B, S2B). For each domain, we resampled the mass scaling curves with centers inside the “domain center region” (Fig. 3C) and determined mass scaling behavior from these “domain centers” up to *r* = 400 nm for A549 cells and *r* = 200 nm for BJ cells. In each individual domain, the mass scaling curve exhibits regimes characterized by a gradual deviation from the initial power-law at larger length scales (Fig. 3D). We performed linear regression on the 2D mass scaling curve and obtained a slope, *D_log_* = 2.71 ± 0.02 for 2 nm < *r* < 102 nm (Fig. 3D, blue dashed line). This power-law scaling relationship can model the mass scaling curve with less than 5% error within the given fitting range, while a more significant divergence is observed beyond *r* = 102 nm (Fig. 3D, red asterisk). Therefore, from the mass scaling curve for a single packing domain, we observe that the smaller length scales have a packing scaling *D* < 3, indicating mass scaling behavior that is not space-filling, and that as *r* increases up to around 100 nm, there is a sharp transition to the supradomain regime with *D_log_*=3. This increase in *D* towards a value of 3 potentially indicates overlap between neighboring domains.

**Fig.3.**
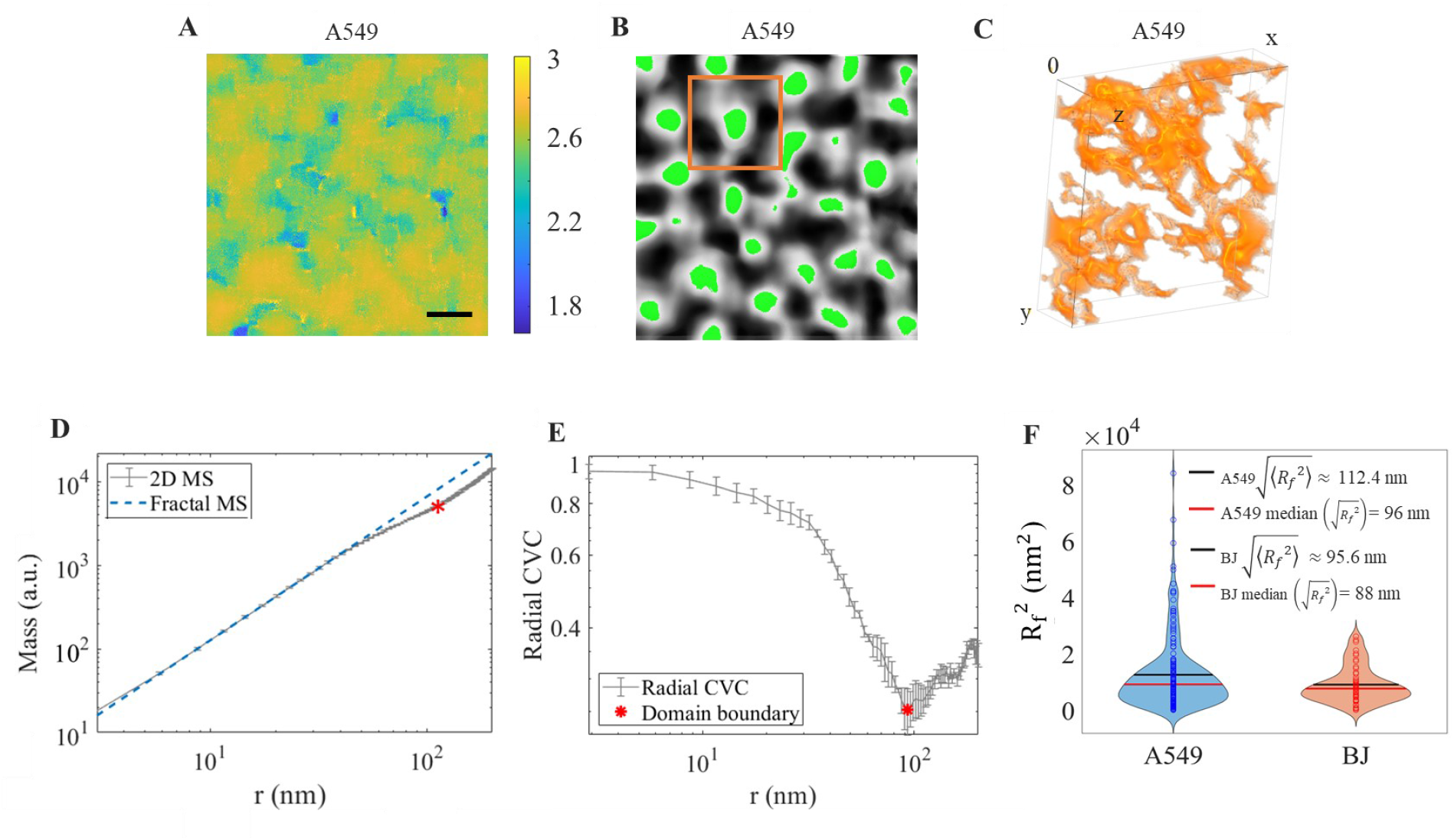
Quantifying domain size and chromatin packing behavior at the domain boundaries. Chromatin packing domains are structurally heterogeneous and anisotropic. (**A**) The spatial mapping of packing scaling *D* within the domain regime for one field of view of an A549 cell. Scale bar: 200 nm. (**B**) From the grayscale representation of **A** after smoothing and local contrast enhancement, the domain centers are identified by the top 10% of the D value. For one domain, the mass scaling curve is resampled from centers within the domain center (green region). (**C**) 3D rendering of the surface of chromatin fibers in a region, including the packing domain of interest (orange square in **B**). (**D**) The average 2D mass scaling (MS) curve of the chromatin within the region of interest (orange square in **B** and **C**). The mass scaling analysis is conducted with randomly selected centers within the domain center. The error bar represents the standard deviation. The MS curve starts with *D* < 3 (blue dashed line) and transitions to values closer to *D* = 3 (beyond the red asterisk). (**E**) Radial distribution of CVC for the same domain shown in **B**, **C**, and **D**. The radial CVC initially slowly decreases within the domain regime. As the length scale approaches the domain boundary (red asterisk), the radial CVC rapidly dips which is followed by a recovery, due to the presence of other domains at those length scales. (**F**) The distribution of *R_f_*^2^, the square of radius of the packing domain, for A549 (blue) and BJ (orange) cells.

Additionally, we calculated the radial distribution of chromatin volume concentration (radial CVC, supplementary methods) to investigate the chromatin packing density from the “domain center region” to the periphery of the individual domain (Fig. 3E). We observed three key trends in the radial CVC at different distances from the domain center: 1) a relatively flat, slowly decreasing curve near the domain center, 2) a rapidly decreasing curve at a moderate distance from the domain center, and 3) an increasing curve at even larger distances. This third trend is likely caused by the inclusion of chromatin from other nearby domains. The transition point from rapid decrease to increase in radial CVC (red asterisk in Figure 3E) is consistent with the transition point in mass scaling from similar mass scaling behavior to random packing (red asterisk in Figure 3D), and both are indicative of the edge of the analyzed domain.

For each domain, we quantified the regime of similar mass scaling behavior, or the radius of the packing domain (*R_f_*), as the smallest length scale that satisfies the following criteria (Fig. S4): (1) Mass scaling curve deviates from the initial power-law calculated from small length scales by 5%, suggesting a significantly different packing behavior; (2) Local packing scaling *D* reaches 3, implying a random structure; (3) The absolute value of the second derivative of the logarithm of the mass scaling curve is greater than 2, indicating a divergence from power-law scaling behavior; (4) The radial CVC starts to increase. We observed a broad range of *R_f_* for both A549 cells with 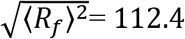 nm and BJ cells with 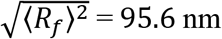 (Fig. 3F). We interpreted *R_f_* as the length scale where the chromatin mass scaling no longer follows a power-law relationship, or where a single packing scaling is not sufficient to explain the packing behavior. However, this view does not indicate each domain is spherical with radius *R_f_*. We further quantified the shape of the domain boundary by calculating the 2D asphericity (*A_s_*) of the chromatin enclosed by the domain boundary (*55, 56*). Considering a 2-dimensional ellipse, 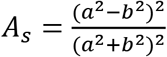, where *a* and *b* are the semiaxes of the ellipse. Here, *A_s_* can take on values from 0 to 1, depending on the ratio (Fig. S4F). For the case *a* = Ď, *A_s_*=0 indicating an isotropic or spherical configuration. In the limit, *a* >> *b*, *A_s_*=1 indicating a linear or stretched configuration. To avoid edge effects, we only considered domains that are entirely within the field of view. We estimated the average of *A_s_* to be 0.4527 ± 0.01 from 134 domains for A549 cells and 0.4511 ± 0.02 from 16 domains for BJ cells, respectively (Fig. S4F). Thus, the packing of chromatin fibers into random domains is both heterogeneous and anisotropic.

### Differential morphological properties of chromatin packing domains

In the context of chemical reactions, macromolecular crowders are any protein, nucleotide, or other macromolecule that occupies physical space but does not directly participate in the reaction (*57, 58*). As transcription reactions are chemical reactions, crowding directly influences both the kinetics and efficiency of transcription (*44*). We have previously developed a computational model of transcription in a realistic chromatin environment by considering chromatin density as the major crowder in the nucleus (*42, 45*). Statistical descriptors of packing domains, including chromatin packing scaling, average chromatin density, and size of domains were determined to be physical regulators of transcription by controlling chromatin density distribution within these domains (*45*). Thus, characterizing the distribution of such properties and understanding their link with chromatin packing can help decode the complex chromatin structure-function relationship (*44*).

First, we determined the average CVC, or chromatin density, per domain to quantify chromatin compaction. Similar to the anisotropy analysis, we excluded the domains at the edge of the field of view. We observed that 95% of 124 domains (110 domains for A549 and 14 domains for BJ) have a CVC within 24% to 49%, with a mean CVC of 36.6% (Fig. 4A). For the same domains, we obtained the distribution of packing scaling *D*, with a mean value of 2.57 ± 0.01 for A549 cells and *D* = 2.65 ± 0.03 for BJ cells (Fig. 4B). For a polymer that exhibits mass scaling behavior within a certain regime, the relationship between CVC and packing scaling follows the relationship 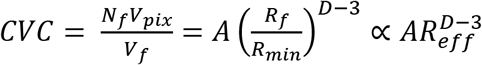, where the total mass of a domain 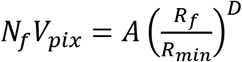 and is measured as product of the number of pixels occupied by chromatin within the domain, *Nf*, and the resolution or smallest unit of chromatin measured by ChromSTEM, *V_pix_*. *R_f_* and *V_f_* are the domain size and total volume, *R_min_* is the radius of the elementary unit of the chromatin chain, *A* is the packing efficiency factor of the fundamental chromatin unit within the domain, and 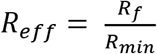 is the effective domain size (*59*). A chromatin polymer with *A* = 1 specifies that each *R_min_* concentric layer of the packing domain is packed in the most efficient manner designated by the domain packing scaling, with the chromatin chain as the primary building block. Here, we assume that the packing efficiency within the elementary unit of the chromatin chain is 1, i.e., the entire volume of the chain is completely filled by chromatin. Similar to *R_f_*, *R_min_* can be estimated from the ring mass scaling curve as the upper bound of the chromatin chain regime, or, in other words, the spatial separation that significantly deviates from the behavior within the chromatin chain (Supplementary Methods). Next, we investigated whether the packing efficiency and the chain size are conserved across a population of isogenic cells or across cell lines with differential genetic makeup by determining the relationship between domain CVC and 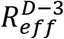. We observed a positive correlation between CVC and *D* for A549 cells (linear regression r^2^ = 0.53) and a weaker correlation for BJ cells (linear regression r^2^ = 0.41) (Fig. 4E). Given that *R_min_* from the mass scaling analysis was not significantly different across domains, cells within the same cell lines, and even between the two cell lines, this relationship suggests that the chain size may be constant across genetically different cells, although the packing efficiency is domain specific. Average *A* for each cell line was evaluated from the regression of CVC on 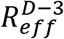. Comparing the A549 and BJ cell lines, we observed a significant difference between the slopes of linear regression (ANCOVA p < 0.0005).

**Fig.4.**
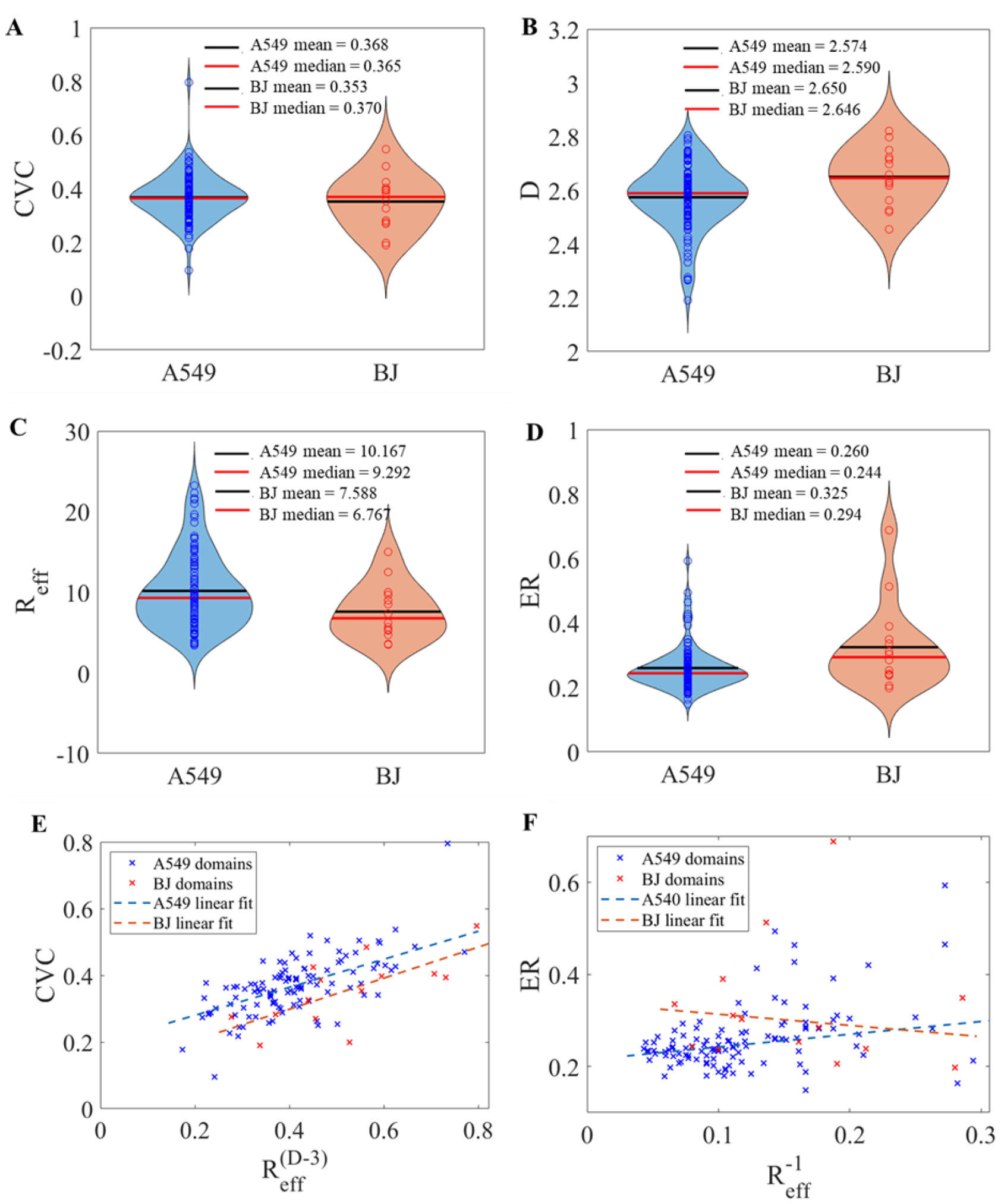
Differential properties of chromatin packing domains. (**A**) Chromatin volume concentration (CVC) distribution per packing domain for A549 cells and BJ cells. A total of 110 A549 cell packing domains and 14 BJ cell packing domains were analyzed. We observed the CVC distribution ranges from 0.10 to 0.80 with a mean value of 0.37 for A549 cells, and the CVC distribution ranges from 0.19 to 0.55 with a mean value of 0.35 for BJ cells. (**B**) For the same domains, we observed packaging scaling *D* varies from 2.19 to 2.80 with a mean equal to 2.57 for A549 cells and *D* from 2.45 to 2.82 with a mean equal to 2.65 for BJ cells. (**C**) Effective domain size *R_eff_* for A549 and BJ cells. The effective domain size is the ratio between domain size *R_f_* and domain chain size *R_min_*. For A549, R_f_ ranges from 3.4 to 23.3 with a mean value of 10.2. For BJ, Rf ranges from 3.5 to 15.0 with a mean of 7.6. (**D**) Exposure Ratio (ER) per domain fraction is defined as the fraction of chromatin voxels on the surface of the interchromatin voids. For A549, the ER ranges from 0.15 to 0.59 with a mean value of 0.26. For BJ, the ER ranges from 0.20 to 0.69 with a mean of 0.33. (**E**) A moderate correlation between domain CVC and *D* has been observed for A549 cells, with r^2^ = 0.53, and a weaker correlation with r^2^ = 0.41 for BJ cells. (**F**) Exposure ratio is positively correlated with inverse effective domain size with the strong linear coefficient for A549, characterized by r^2^ = 0.66, but showed a very weak negative correlation for BJ, with r^2^ = 0.33.

As the boundaries of TADs and chromatin domains are enriched in active transcription processes (*59, 60*), we next studied how the probability of chromatin elements being exposed on the surface of domains changes across domains and across cell lines. We defined an exposure ratio (*ER*) as the fraction of ChromSTEM voxels on the surface of the domain compared to the total number of pixels encompassing the domain volume. The surface here exclusively refers to the internal surface created by the interchromatin voids within domains. This metric can be thought of as a way of evaluating the surface area to volume ratio of a domain. Without changing the genomic size of a domain, an increase in *ER* for a given chromatin domain would indicate an increase in the chromatin domain surface, which could increase the amount of surface chromatin that is accessible to transcription processes. First, we define *A_sp_* as the surface packing efficiency, i.e. the prefactor in the scaling relationship 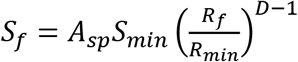 where *S_f_* is the total intradomain surface area and *S_min_* is the surface area of the elementary unit of the chromatin chain, measured as the number of pixels. For a domain, *ER* can be then estimated by the following: 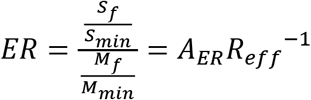, where 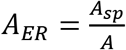 is the exposure ratio efficiency factor and represents the ratio between the packing efficiency at the surface compared to throughout the entire domain. Next, we investigated if *A_ER_* is constant for all domains within the same cell line and across different cell lines. We calculated the ER for each domain and the corresponding inverse effective domain size, *R_eff_*. Although the distribution of *R_eff_* is relatively similar between the cell lines (Fig. 4C), the variability of exposure ratios within each cell line is markedly different (Fig. 4D). Next, we performed linear regression analysis to better characterize the relationship between the inverse effective domain size and ER at the domain level. We observed a strong positive association between the *ER* and *R_eff_*^-1^ for A549 cells (linear regression r^2^ = 0.66) and a weak, negative correlation for BJ cells (linear regression r^2^ = 0.33) (Fig. 4F). This suggests that for A549 cells, *A_ER_* varies around a common average value but is not universal for all domains. On the other hand, the incongruent trend for the BJ cells suggests that *A_ER_* might be significantly different across cell lines. Altogether, these results demonstrate that domains have transcriptionally-relevant properties, including average density, packing scaling, packing efficiency, and exposure ratios, that are heterogeneous within the same cell line, but the differences across cell lines is even larger.

## Discussion

Utilizing ChromEM staining that selectively enhances the contrast of DNA and dual-axis electron tomography with high-angle annular dark-field imaging mode, ChromSTEM has the advantage of resolving chromatin packing in 3D at a sub-3-nm spatial resolution at the single-cell level (Fig.1). Employing ChromSTEM on two genetically different cell lines, both chemically fixed A549 cells (cancer) and BJ cells (non-cancer), we were able to quantify chromatin packing *in vitro* down to the level of the primary chromatin fiber. Importantly, we studied these cell lines to distinguish basic principles behind chromatin packing that are generally cell line-invariant and cell line-specific. We do not assume that the exact results from A549 cells extend to all cancerous cells and the results obtained from BJ cells represent all non-cancerous cells. By analyzing the mass-scaling behavior of the chromatin polymer, we observed spatially separable and geometrically anisotropic packing domains ~100 nm in radius in both cell lines (Fig. 2&3). The mass scaling within the packing domains follows a power-law relationship with *D*<3, indicating that chromatin packs into higher-order domains with similar mass-scaling behavior, and that packing domains have radially arranged layers with decreasing chromatin density from the domain center to the periphery. This “core-shell” structure supports earlier experimental work using super-resolution microscopy at a coarser spatial resolution (*57*).

Although they have random arrangements in three-dimensional space, these domains are intrinsically not space-filling. This morphology is also validated by the fact that CVC values are always smaller than 100%. In addition, the local density progressively decreases from the domain center region to the periphery, which agrees with findings using other microscopic studies (*61*). At the same time, the domains are not completely isolated from each other without any chromatin density in between, as CVC values are always above 0. From these observations, it is reasonable to suggest chromatin is organized into complex, porous packing domain structures which are connected by less dense chromatin fibers. The porosity of domains could provide additional surface area, potentially promoting diffusion and targeted search mechanisms, such as transcription. Outside of packing domains, the packing scaling increases to 3 after crossing the domain boundary, potentially indicating a random distribution of multiple domains with respect to each other.

Interestingly, the previous ChromEMT study did not observe any higher-order chromatin structures above the level of the primary fiber (*15*), which is incongruous with other EM and optical microscopy studies. The size of the packing domains observed using ChromSTEM (~200nm diameter) are consistent with previous observations of higher-order chromatin domains, including ‘chromomeres’ (~200-300nm) (*21*), replication domains (~110-160nm)(*22–25*), and domains associated with TADs (~200-300nm)(*62*), although future experiments are necessary to elucidate the molecular basis of these packing domains. Importantly, ChromSTEM imaging and analysis were able to characterize the structural properties of these domains with high fidelity.

Our previous experiments on isogenic cell lines have demonstrated *D* as a crucial modulator of transcriptional plasticity (*45*). However, in these experiments either the expression levels of a certain chromatin-associated gene were altered, or a cell population was treated with a *D*-lowering agent, both while keeping other transcriptional regulators fairly similar. In general, cell lines from cancer patients which have been immortalized and propagated over generations cannot be considered genetically similar to normal, non-cancer human cells. Thus, this comparison cannot be directly extended to cell lines with distinct germline profiles such as A549 (cancer) and BJ (non-cancer). In this study, we obtained the distribution of packing scaling *D*, with mean values of *2.57* ± 0.01 for A549 cells and *D* = 2.65 ± 0.03 for BJ cells (Fig. 4B). In this case, other factors in addition to *D* may be playing a role in influencing the transcriptional plasticity of these two genetically distinct cell lines, such as the alteration in the availability of transcriptional machinery, making a direct comparison extremely convoluted (*44, 45*). Furthermore, for packing domains of both A549 and BJ cell lines, we observed a diverse range of CVC, *R_f_*, *A_s_*, and *ER* values, all of which could also impact transcription rate (Fig. 4). As domains adopt a packing structure with similar mass scaling behavior, some of these morphological properties are interrelated from a polymer physics perspective. From ChromSTEM data, we observed chromatin density and packing scaling cannot be related by a universal factor, and this packing efficiency varies depending on domain size and cell type. We also observed a similar complex relationship between the exposure ratio, the probability of a chromatin segment to be on the domain surface, and domain size, which is not universal for all domains within a cell line and varies significantly across cell lines. The differential properties of domains could potentially play a role in regulating gene activities by controlling the size of proteins and other macromolecular complexes that can navigate through this network, thus influencing material transportation and gene accessibility.

The major limitations of ChromSTEM include chemical fixation, low throughput due to electron tomography, and inability to obtain loci-specific gene accessibility information. Therefore, ChromSTEM findings are not directly comparable to discoveries made from sequencing-based techniques such as Hi-C or loci-based imaging methods such as Fluorescence In Situ Hybridization (FISH). Despite the limitations, we believe that ChromSTEM and the associated analysis methods developed in this work should become an important tool for the understanding of the 3D structure of chromatin and its function. Additionally, we have recently demonstrated that the Nanoscale Chromatin Imaging and Analysis (nano-ChIA) platform allows quantification of different aspects of the chromatin structure by combining high-resolution imaging using ChromSTEM with other nanoimaging techniques involving labeling of molecular functionality and high-throughput chromatin dynamics imaging in live cells. In the future, co-registering ChromSTEM with 3D super-resolution techniques labelling markers for heterochromatin and euchromatin (*31*) will be integral to improving our understanding of the relationships between the physical structure of chromatin within packing domains, epigenetic modifications and transcription. Future work should also focus on developing novel labeling methods that target particular genes that are compatible with ChromSTEM sample preparation and imaging, and colocalizing chromatin morphological and genetic information for a greater number of cells (*58*).

## Materials and Methods

### Cell culture

A549 cells were cultured in Dulbecco’s Modified Eagle Medium (ThermoFisher Scientific, Waltham, MA, #11965092). BJ cells were cultured in Minimum Essential Media (ThermoFisher Scientific, Waltham, MA, #11095080. All cells were maintained at physiological conditions (5% CO2 and 37 °C). Experiments were performed on cells from passages 5–20.

### ChromEM sample preparation

The ChromSTEM sample staining and resin-embedding followed the published protocol (*15*), and detailed reagents and steps can be found in **Table S1**. All cells were thoroughly rinsed in Hank’s balanced salt solution without calcium and magnesium (EMS) before fixation with EM fixative. Two stages of fixation were performed: room temperature fixation for 5 min and on-ice fixation for an hour with fresh fixative. The cells were kept cold for all following steps before resin embedding either on ice or a cold stage with the temperature monitored to vary from 4°C to 10°C. The biopsy of the mouse ovary was embedded in low melting point agarose (Thermo Fisher) and 40 μm thick sections were prepared using a vibratome (VT1200 S Leica) on ice. The sections were deposited onto a glass-bottom petri-dish (MatTek) and treated as described for cells in the following steps.

After fixation, the samples were bathed in blocking buffer for 15 min before being stained by DRAQ5^TM^ (Thermo Fisher) for 10 min. The cells were rinsed and kept in the blocking buffer before photo-bleaching and submerged in 3-5’-diaminobenzidine (DAB) solution (Sigma Aldrich) during photo-bleaching on the cold stage.

A Nikon microscope (Nikon Inc.) was used for photo-bleaching. A cold stage was developed in-house from a wet chamber equipped with humidity and temperature control. After photo-bleaching, the cells were rinsed in 0.1M sodium cacodylate buffer thoroughly. Reduced osmium solution (EMS) was used to enhance the contrast in STEM HAADF mode, and the heavy metal staining lasted 30 min on ice. Serial ethanol dehydration was performed, and during the last 100% ethanol wash, the cells were brought back to room temperature. Durcupan resin (EMS) was used for embedding after infiltration, and the blocks were cured at 60°C for 48 hrs.

An ultramicrotome (UC7, Leica) and 35 degree diamond knife (DiATOME) were employed to prepare sections of different thicknesses. For STEM HAADF tomography, semi-thick sections were made and deposited onto a copper slot grid with carbon/Formvar film. All TEM grids were plasma cleaned before sectioning and no post-staining was performed on the sections. 10 nm colloidal gold fiducial markers were deposited on both sides of the sample.

A step-by-step protocol can be found in **Protocol S1**.

### EM data collection and tomography reconstruction

A 200kV cFEG STEM (HD2300, HITACHI) with HAADF mode was employed for all image collection. For tomography, the sample was tilted from −60° to 60° with 2° increments on two roughly perpendicular axes. Each tilt series was aligned with fiducial markers in IMOD and reconstructed using Tomopy (*46, 47*) with a penalized maximum likelihood for 40 iterations independently. IMOD was used to combine the tomograms to suppress artifacts (Fig. S5), the nominal voxel size varies from 1.8 nm to 2.9 nm for different samples (*47*). Volume Viewer in FIJI was employed for surface rendering (*15*).

### Chromatin mask segmentation and mass scaling analysis

We generated binary masks for chromatin from the ChromSTEM tomograms based on automatic thresholding in FIJI as reported previously with fine-tuned imaging processing parameters. For all chromatin masks used in this work, the following procedure was performed. First, the local contrast of the tomograms was enhanced by CLAHE, with a block size of 120 pixels. Then, Ostu’s segmentation algorithm with an automatic threshold was employed. Finally, we removed both dark and bright outliers using a threshold of 50 and a radius of 2 to refine the chromatin mask.

In polymer physics, mass-scaling is the relationship between the material *M* within concentric circles of radius r. For a polymer with power-law mass scaling behavior, the mass scales as *M*(*r*) ∝ *r^D^*, where *D* is the power-scaling exponent or fractal dimension. In the 1D scenario, *M* is the amount of chromatin positioned on the perimeter of a circle (*P* = 2*πr*). We referred to the 1D case as “ring mass scaling”. For 2D, *M* is the amount of chromatin enclosed by a circle with area *A* = *πr*^2^; for 3D, *M* is the amount of chromatin within a sphere with volume 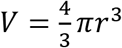. To calculate mass scaling, starting from the binarized chromatin mask, concentric circles with radius from 1 pixel to 300 pixels were employed with non-zero centers were randomly selected within the sample, and *M* is calculated as the total number of non-zero voxels within the predefined space (perimeter, area, and volume). For each stack of tomography, we averaged mass scaling curves with different centers at each dimension. For A549 cells, 1D, 2D, and 3D mass scaling were analyzed. For BJ cells, only 1D and 2D mass scaling were evaluated, as the thickness of the tomography is limited. We found from our calculations from polymer trajectories that the 3D mass scaling exponent can be approximated using the 2D case and the 1D case (Fig. S3A): *D*_3*D*_ = *D*_2*D*_ + 1, and *D*_3*D*_ = *D*_1*D*_ + 2, with standard errors of the mean of 0.023 and 0.019 respectively.

### Domain and boundary analysis

Unlike the mass scaling for the entire sample where we averaged randomly sample mass scaling curves from different regions mentioned above, to identify domains, we used the local relationship between mass scaling curves and *r*. To obtain a smooth mapping of the packing scaling, we calculated the mass scaling within a moving window (300 pixels x 300 pixels) with a stride of 1 pixel on each virtual 2D tomogram. To remove noise at small length scales, we sampled all the non-zero pixels as the center of the mass scaling analysis within the center of the window (10 pixels by 10 pixels) and obtained the average. We then calculated the packing scaling *D* from the averaged mass scaling curve per window by linear regression, then mapped the value to the center of the window.

The above method provided the spatial distribution of packing scaling, though it is capable of identifying domains and locating the domain center region, it is not sufficient to isolate domain boundary precisely nor provide accurate values for each domain, as the moving window approach is equivalent to performing filtering. To obtain domain boundary, we calculated the mass scaling again but centered within each domain center.

The domain center is calculated from the *D* map from the moving average method (Fig. S2B). The starting point of the analysis is the *D* map calculated by the moving window mass-scaling in grayscale. Then we applied Gaussian filtering with radius = 5 pixels followed by CLAHE contrast enhancement with a block size of 120 pixels in FIJI (*15*). With a “flooding” algorithm in the MATLAB image segmentation GUI, we identified the center of the domains (green) at automatic thresholding values and created the binary mask for those regions accordingly. We then identified the center pixel of gravity per binary domain center. To obtain the mass scaling curve for a single domain, we first sampled multiple mass scaling curves with centers on the nonzero pixels around the center pixel within a 10-pixel x 10-pixel window. We then used the average mass scaling curve for that domain.

To obtain the domain boundary, we utilized the mass scaling behavior of the packing from the center region to the periphery. To evaluate such behavior, besides the mass scaling curve, we also leveraged the radial volume chromatin concentration (CVC). We adopt the definition of CVC from published work (*15*): which is the fraction of volume occupied by chromatin. This value is calculated from the binary chromatin mask obtained from tomography data. The boundary of the domain can be seen as the length scale where a single power-law relationship no longer holds is defined as the domain size *R_f_*. Practically, we used the smallest length scale that meets at least one of the four criteria: 1. Mass scaling curve deviates from the initial power-law calculated from small length scales by 5%, suggesting a significantly different packing behavior; 2. Local packing scaling *D* reaches 3, implying a random structure; 3. The absolute value of the second derivative of the logarithm of the mass scaling curve is greater than 2, indicating a divergence from the power-law. 4. The radial CVC starts to increase. An example of the workflow to determine *R_f_* can be found in Fig. S4. Similar to *R_f_*, the average size of the fundamental building block of the domain *R_min_* can be measured by the spatial separation where the mass scaling behavior deviates significantly (5%) from the behavior within the initial chromatin chain regime. In this work, the initial mass scaling behavior is quantified by the slope of the first two data points on the 2D mass scaling curve. The ratio between *R_f_* and *R_min_* is defined as the effective domain size *R_eff_*.

### Domain morphological property analysis

We calculated four different morphological properties for each domain: packing scaling *D*, asphericity *A_s_*, domain CVC, and exposure ratio (ER). Chromatin packing scaling as *D_log_* was estimated by the slope of the linear regression of the average mass scaling curve in log-log, fitted from r ~10 nm to r~30 nm. Chromatin within *R_f_* distance from the domain center pixel was selected as a “domain”, though realistically the domains are likely to adopt an irregular shape. The 2D asphericity *A_s_* is calculated slice-by-slice using the following expression: 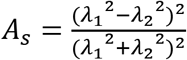, where *λ*_1_ and *λ*_2_ are the eigenvalues of the 2D gyration tensor of the domain (*55*). Then we calculated the mean value from each slice to be the *A_s_* for that domain. The CVC is calculated as the ratio of the total number of nonzero (chromatin) voxels over the total number of voxels per domain. And the exposure ratio is the fraction of voxels on the domain surface. The surface of the domain includes only the surface of the pores within the domain, it excludes the external surface created by an artificial boundary imposed by *R_f_*.

## Acknowledgments

This work was supported by Medical Scientist Training Program T32GM008152, National Institutes of Health grants U54CA193419, R01 CA228272, R01CA225002, and National Science Foundation grants EFMA-1830961 and EFMA-1830969. This work made use of the BioCryo facility of Northwestern University’s NUANCE Center, which has received support from the SHyNE Resource (NSF ECCS-2025633), the International Institute for Nanotechnology (IIN), and Northwestern’s MRSEC program (NSF DMR-1720139). The imaging methodology is partially based on research sponsored by the Air Force Research laboratory under agreement number is FA8650-15-2-5518. The U.S. Government is authorized to reproduce and distribute reprints for Governmental purposes notwithstanding any copyright notation thereon. The views and conclusions contained herein are those of the authors and should not be interpreted as necessarily representing the official policies or endorsements, either expressed or implied, of Air Force Research Laboratory or the U.S. Government.

## Author Contributions

Y.L., V.P.D., and V.B. conceived the study, Y.L. and V.A. performed the ChromSTEM experiments. Y.L., V.A., R.K.V, and W.L. analyzed the data. A.E. and J.F. assisted with the ChromSTEM experiments. E.R. and R.B. participated in the sectioning of ChromSTEM samples. K.H., L.A., and M.C. participated in the data analysis. I.S., V.P.D., and V.B. supervised the project. All authors reviewed and edited the manuscript.

