## Supplemental Information for "Statistical Analysis of Three-Dimensional Chromatin Packing Domains Determined by Chromatin Scanning Transmission Electron Microscopy (ChromSTEM)"

**Supporting Information**

**Mov. S1.** Tomogram of the 3D chromatin structure moving through z-direction of the chromatin in the A549 cell in Fig. 1

**Mov. S2.** Tomogram of the 3D chromatin structure moving through z-direction of the chromatin in the BJ cell in Fig. S1

**Mov. S3.** Tomogram of the 3D chromatin structure moving through z-direction of the chromatin in another A549 cell

**Mov. S4.** 3D rendering of the entire tomography stack of **Mov. S1**

**Mov. S5.** 3D rendering of nucleosomal superstructures from the same A549 cell as in **Mov. S1**

**Mov. S6.** 3D rendering of the entire tomography stack of **Mov. S2**

**Mov. S7.** 3D rendering of nucleosomal superstructures from the same BJ cell as in **Mov. S2**

**Mov. S8.** 3D rendering of the entire tomography stack of **Mov. S3**

**Mov. S9.** 3D rendering of nucleosomal superstructures from the same A549 cell as in **Mov. S3**

**Protocol S1. Sample preparation for ChromSTEM for cell cultures**

**Fixation:**

1. Wash the cells in the petri-dish in the washing solution for 3 times, 2 minutes each.
2. Fix the cells with the fixation solution for 5 minutes at room temperature.
3. Continue to fix the cells with fresh fixation solution for an additional 1 hour on ice.

The following steps before the last ethanol dehydration are either on ice or a cold stage, all reagents must be chilled to 4$^{o}$C before use.

**DNA Staining:**

4. Wash the cells with 0.1M sodium cacodylate buffer for 5 times on the ice, 2 minutes each.
5. Block the cells with a blocking solution for 15 minutes.
6. Stain the cells with DNA staining solution for 10 minutes.
7. Wash the cells with the blocking solution 3 times, 5 minutes each.

**Photo-bleaching:**

8. Bath the cells in the bathing solution before photo-bleaching
9. Photo-bleach the cells using continuous epi-fluorescence illumination (150 W Xenon Lamp) with Cy5 red tilter and a 100x objective for 7 minutes for each spot on the cold stage.
10. Replace the bathing solution in the petri-dish with a fresh bathing solution every 15 minutes (roughly two spots).

**Heavy metal staining:**

11. Rinse the cells with 0.1 M sodium cacodylate buffer 5 times, 2 minutes each.
12. Stain the cells with reduced osmium staining solution for 30 minutes.
13. Wash the cells with double distilled water 5 times, 2 minutes each.

**Dehydration and Resin embedding:**

14. Dehydrate the cells with serial ethanol (30%, 50%, 70%, 85%, 95%, 100% twice) on ice, 2 minutes each.
15. Wash the cells with 100% ethanol at room temperature for 2 minutes.
16. Infiltrate the cells with a 1:1 infiltration mixture at room temperature for 30 minutes.
17. Infiltrate the cells with a 2:1 infiltration mixture at room temperature for 2 hours.
18. Infiltrate the cells with Durcupan$^{TM}$ resin mixture 1 at room temperature for 1 hour.
19. Infiltrate the cells with Durcupan$^{TM}$ resin mixture 2 at 50$^{o}$C in the dry oven for 1 hour.

Flat embed the cells with fresh Durcupan$^{TM}$ resin mixture 2 in Beem capsule and cure at 60 $^{o}$C in the dry oven for 48 hours.

**Table S1. Reagents used in ChromSTEM Staining**

| Reagent | Formula |
|---|---|
| Washing solution | Hank's balanced salt solution without calcium and magnesium |
| Fixation solution | 2.5% EM grade glutaraldehyde<br>2% paraformaldehyde<br>2 mM $CaCl_2$<br>0.1 M sodium cacodylate buffer, pH = 7.4 |
| Blocking solution | 10 mM glycine<br>10 mM potassium cyanide<br>0.1 M sodium cacodylate buffer, pH = 7.4 |
| DNA staining solution | 10 µM DRAQ5<br>0.1% SAPONIN<br>0.1 M sodium cacodylate buffer, pH = 7.4 |
| Bathing solution | 2.5 mM 3,3'- diaminobenzidine tetrahydrochloride (DAB)<br>0.1 M sodium cacodylate buffer, pH = 7.4 |
| Reduced osmium staining solution | 2% osmium tetroxide<br>1.5% potassium ferrocyanide<br>2 mM $CaCl_2$<br>0.15 M sodium cacodylate buffer, pH = 7.4 |
| Durcupan ™ resin mixture 1 | 10 mL Durcupan ™ ACM single component A, M, epoxy resin<br>10 mL Durcupan ™ ACM single component B, hardener 964<br>0.15 mL Durcupan ™ ACM single component D |
| Durcupan ™ resin mixture 2 | 10 mL Durcupan ™ ACM single component A, M, epoxy resin<br>10 mL Durcupan ™ ACM single component B, hardener 964<br>0.2 mL Durcupan ™ ACM, single component C, accelerator 960<br>0.15 mL Durcupan ™ ACM single component D |
| 1:1 infiltration mixture | 10 mL 100% ethanol<br>10 mL Durcupan ™ resin mixture 1 |
| 2:1 infiltration mixture | 5 mL 100% ethanol<br>10 mL Durcupan ™ resin mixture 1 |

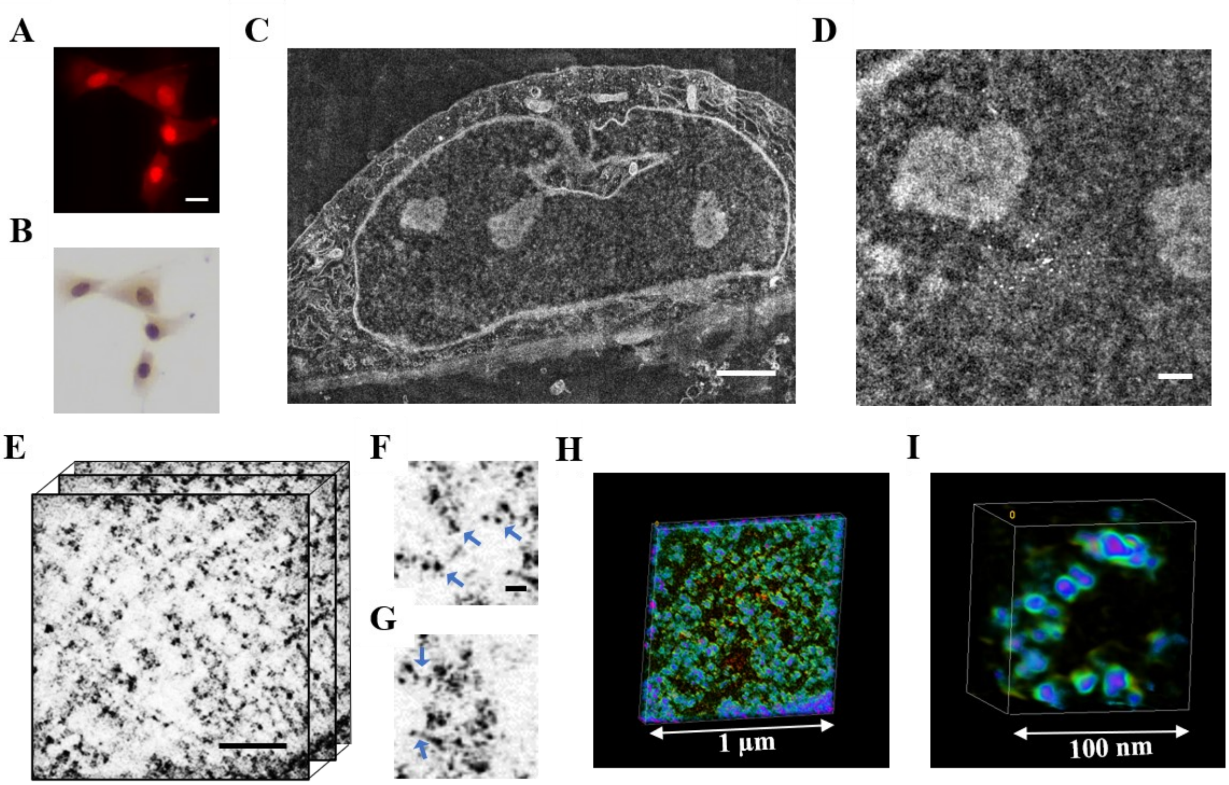


**Fig. S1 ChromSTEM tomography reconstruction of the chromatin of a BJ cell.** (**A**) Photo-oxidation by DRAQ5, fluorescence image of a group of four BJ cells. Scale bar: 20 µm (**B**) Same group of cells after resin embedding show higher contrast under a bright-field optical microscope. (**C**) HAADF image of a BJ cell nucleus prepared with ChromEM staining method. Scale bar: 2 µm. (**D**) HAADF image at higher magnification of the same BJ cell nucleus in (**C**), the colloidal gold nanoparticles are visible and used as fiducial markers in tomography collection. Scale bar: 500 nm. (**E**) A stack of virtual 2D tomograms of the chromatin of the same BJ cell. Scale bar: 200 nm. (**F**-**G**) Magnified views of the tomograms with details of nucleosomal structures (blue arrows in **F**) and linkers between them (blue arrows in **G**). Scale bar: 30 nm. (**H**-**I**) 3D volume rendering, color-coded by the tomogram voxel intensity. The tomogram voxel intensity increases from green to blue to pink. Nucleosomal superstructures with higher voxel intensity are identifiable at higher magnification (**I**).

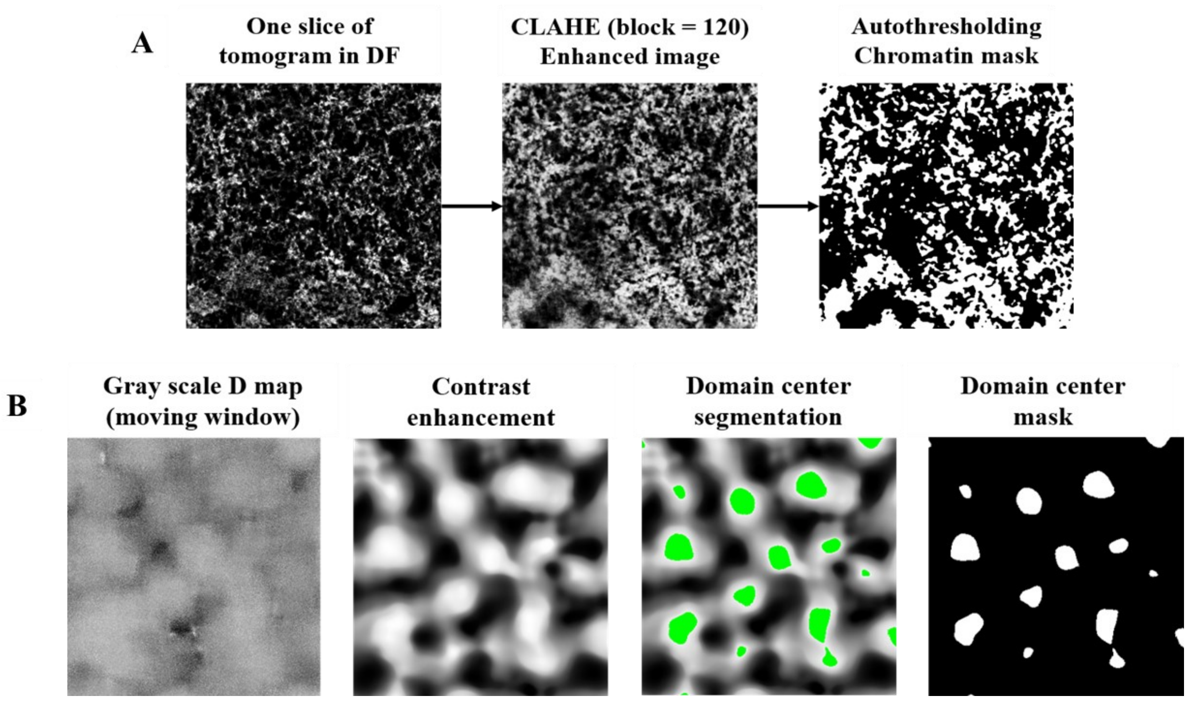


**Fig. S2 Segmentation method to obtain chromatin and domain center masks.**
**(A)** Segmenting tomograms to chromatin mask**:** The segmentation was performed in FIJI. For each frame, we applied local contrast enhancement (CLAHE) with a block size of 120 pixels. We then applied automatic grayscale thresholding to the entire stack of tomograms using Otsu's algorithm. Finally, we polished the mask by removing both dark and bright outliers using a threshold of 50 and a radius of 2. The choice of the block size of CLAHE was optimized to obtain the best segmentation result benchmarked to manual segmentation. **(B)** Domain center region segmentation: The starting point of the analysis is the D map calculated by the moving window mass-scaling in grayscale. Then we applied Gaussian filtering with radius = 5 pixels followed by CLAHE contrast enhancement with a block size of 120 pixels in FIJI. With a "flooding" algorithm in the MATLAB image segmentation GUI, we identified the center of the domains (green) at automatic thresholding values and created the binary mask for those regions accordingly. We then identified the center pixel of gravity per binary domain center. To obtain the mass scaling curve for a single domain, we first sampled multiple mass scaling curves with centers on the nonzero pixels around the center pixel within a 10-pixel x 10-pixel window. We then used the average mass scaling curve for that domain for subsequent analysis.

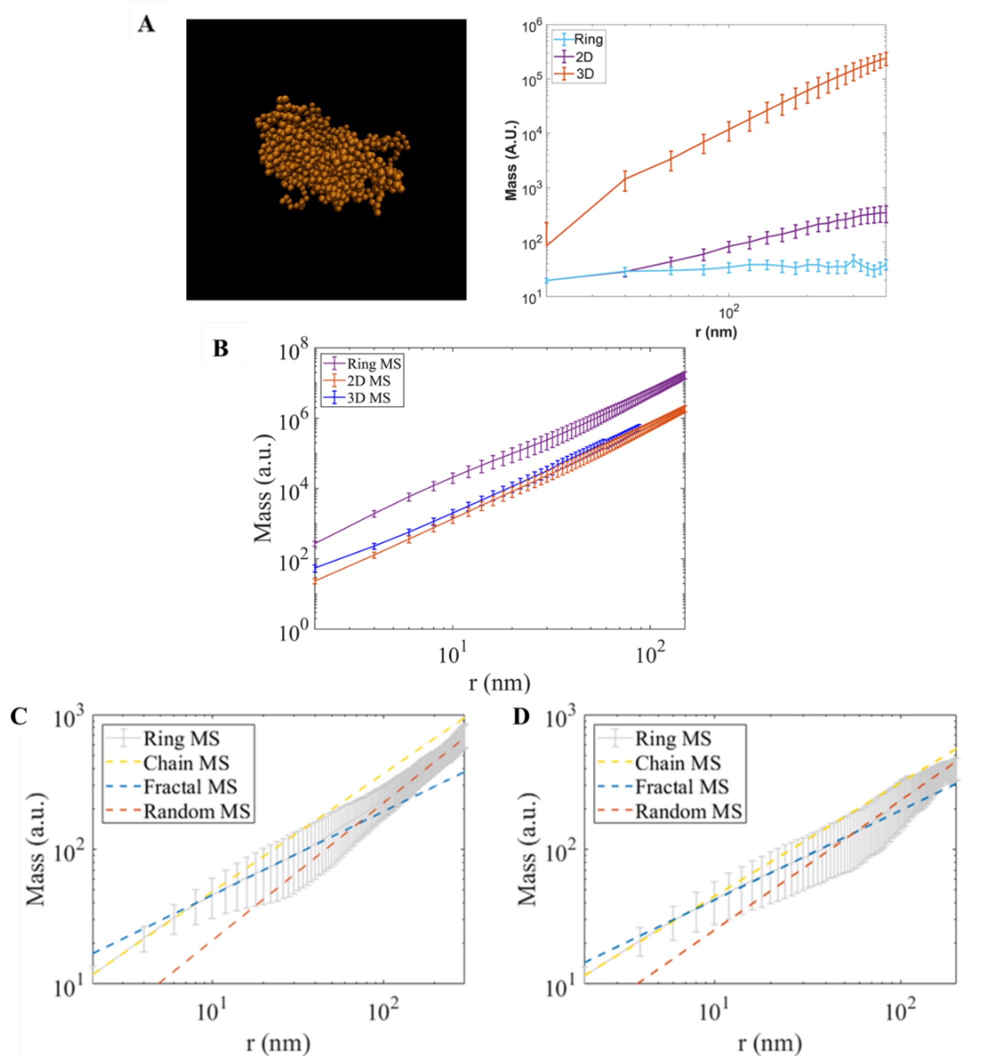


**Fig. S3 Mass scaling analysis at different dimensions.** **(A)** Rendering of a self-attracting homopolymer with $D$ = 2.37 (left) which was estimated from 3D, 2D, and Ring Mass scaling (right) by performing linear regression in the log-log scale on the mass scaling curves for the given dimensions within 20 nm - 240 nm. 3D mass scaling exponent can be approximated using the 2D case and the 1D case: $D_{3D} = D_{2D} + 1$, and $D_{3D} = D_{1D} + 2$, with standard errors of the mean of 0.023 and 0.019 respectively. **(B)** Mass Scaling curves for A549 cells plotted for different dimensions as *3D MS, 2D MS + r, Ring MS + 2r* in the log-log scale. The equivalent slope for the three dimensions in the linear region extending from 2 nm to ~90 nm indicates that the 3D mass scaling exponent can be derived from 2D and Ring Mass scaling exponents. (**C**) The ring mass scaling curve for A549 cells seems to show three regimes: 1. Chain MS with slope, $D = 2.85 \pm 0.07$ fitted from r = 2 nm to 10 nm (yellow dashed line); 2. Domain MS with slope, $D = 2.69 \pm 0.02$ fitted from r = 10 nm to r = 80 nm (blue dashed line); 3. Random MS with slope, $D = 3.02 \pm 0.02$ fitted from r = 140 nm to 160 nm (red dashed line). (**D**) The ring mass scaling curve for BJ cells shows similar three power-law regimes: 1. Chain MS with slope, $D = 2.82 \pm 0.08$ fitted from r = 2 nm to 10 nm (yellow dashed line); 2. Domain MS with slope, $D = 2.67 \pm 0.02$ fitted from r = 10 nm to r = 80 nm (blue dashed line) ; 3. Random MS with slope, $D = 2.98 \pm 0.07$ fitted from r = 95 nm to 120 nm (red dashed line).

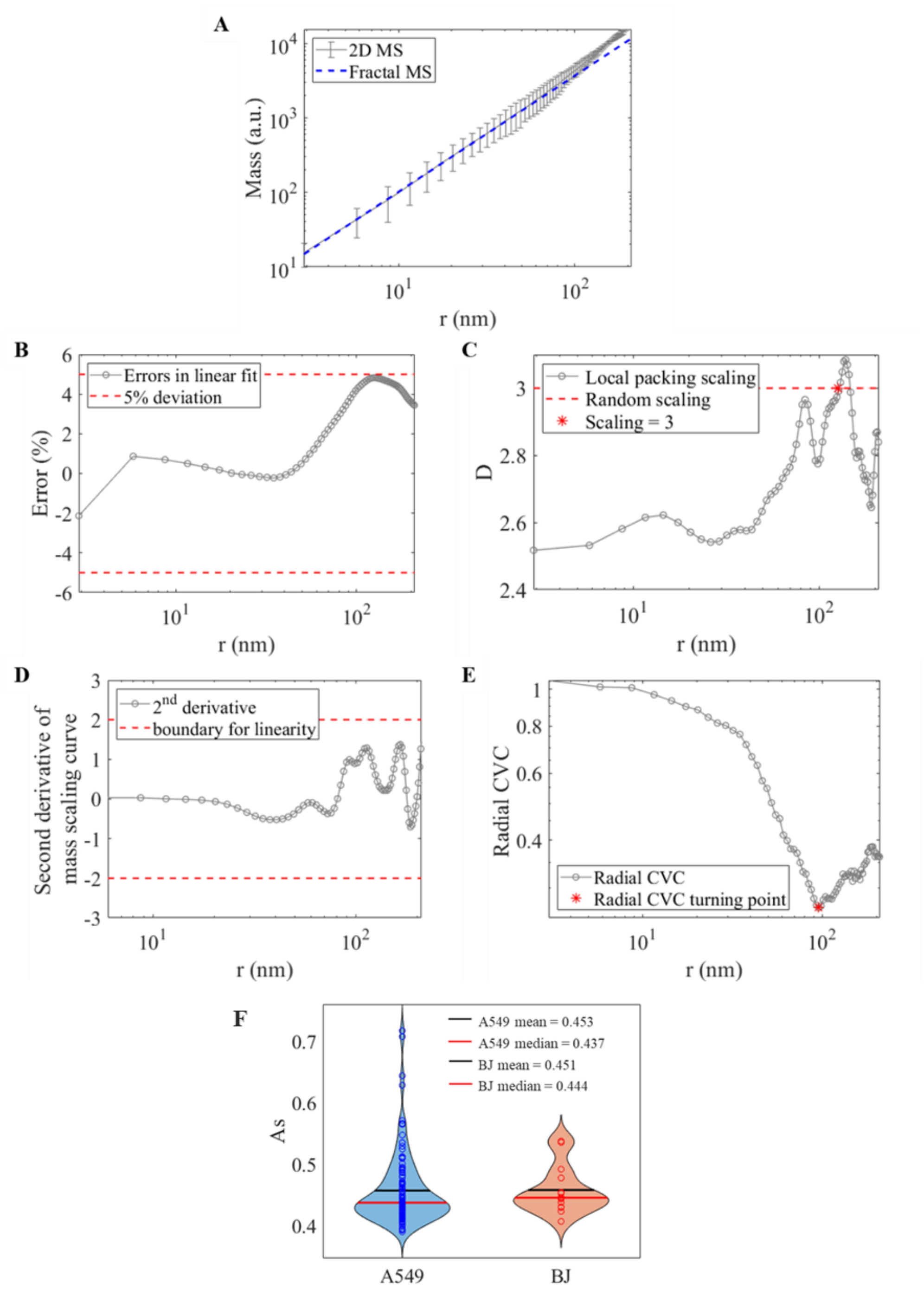


**Fig. S4 Determining domain boundary from the mass scaling behavior.** (**A**) 2D mass scaling curve of one domain. Linear regression was performed from r = 2 nm to r = 100 nm. We obtained a packing scaling of D = 2.7. Beyond r = 100 nm, the mass scaling curve deviates from a power-law mass scaling. We performed the four types of analysis (**B**-**E**) to determine the boundary of domains. (**B**) Mass scaling curve deviates from the initial power-law mass scaling calculated from small length scales by 5%. (**C**) Local packing scaling *D* reaches 3, implying a random structure. Here, the packing scaling = 3 at r = 102 nm. (**D**) The absolute value of the second derivative of the logarithm of the mass scaling curve is greater than 2, indicating a divergence from the power law. Here, all length scales follow under this error margin. (**E**) The radial CVC starts to increase. The radial CVC decreases initially then increases at r = 95.7 nm. We choose the smaller r if they exist. Therefore, in this case, comparing (**C**) and (**E**), we determined the domain size $R_f$ = 95.7 nm. (**F**) The distribution of $A_s$, the asphericity of the chromatin fibers within the domain, for A549 (blue) and BJ (orange) cells.

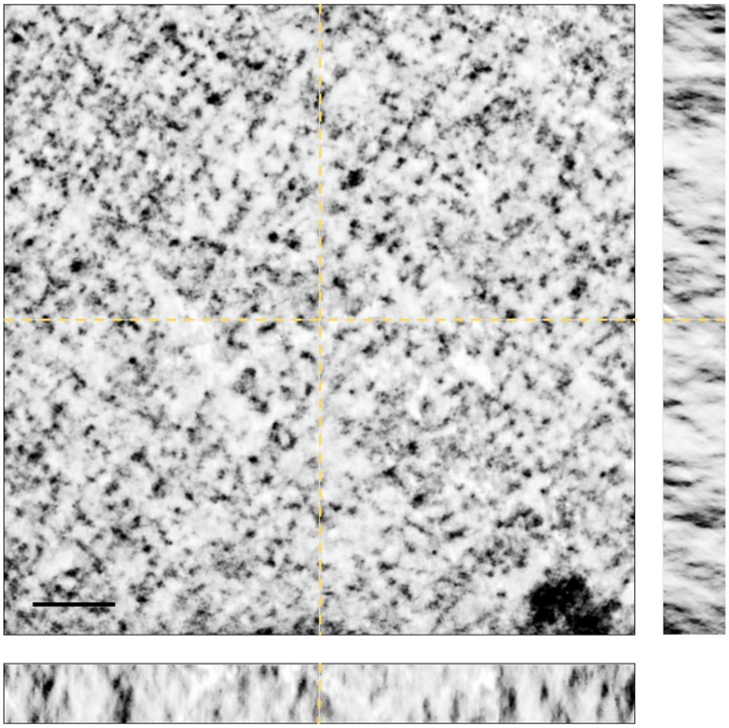

**Fig. S5 Orthogonal view of the reconstructed tomography for A549 chromatin.** To suppress artifacts associated with “missing wedge”, we performed dual-tilt tomography reconstruction in combination with a penalized maximum likelihood reconstruction algorithm. In the side view (x-z and y-z direction, there are still streaks visible due to the “missing cone”, but the contrast of nucleosomes is significantly higher than the noise and sufficient for chromatin mask segmentation. Scale bar: 200 nm.
